# High-quality assembly of sweet basil genome

**DOI:** 10.1101/476044

**Authors:** Nativ Dudai, Marie-Jeanne Carp, Renana Milavski, David Chaimovitsh, Alona Shachter, Kobi Baruch, Gil Ronen, Itay Gonda

## Abstract

Sweet basil, sometimes called the King of Herbs, is well known for its culinary uses, especially in the Italian sauce ‘Pesto’. It is also used in traditional medicine, as a source for essential oils and as an ornamental plant. So far, basil was bred by classical and traditional methods due to lack of a reference genome that will allow optimized application of the most up-to-date sequencing techniques. Here, we report on the first completion of the sweet basil genome of the cultivar ‘Perrie’, a fresh-cut Genovese-type basil, using several next generation sequencing platforms followed by genome assembly with NRGENE’s DeNovoMAGIC assembly tool. We determined that the genome size of sweet basil is 2.13 Gbp and assembled it into 12,212 scaffolds. The high-quality of the assembly is reflected in that more than 90% of the assembly size is composed of only 107 scaffolds. An independent analysis of single copy orthologues genes showed a 93% completeness which reveal also that 74% of them were duplicated, indicating that the sweet basil is a tetraploid organism. A reference genome of sweet basil will enable to develop precise molecular markers for various agricultural important traits such as disease resistance and tolerance to various environmental conditions. We will gain a better understanding of the underlying mechanisms of various metabolic processes such as aroma production and pigment accumulation. Finally, it will save time and money for basil breeders and scientists and ensure higher throughput and robustness in future studies.

## Introduction

Basil, *Ocimum basilicum* L., an important aromatic plant of the *Lamiaceae* family, is grown worldwide both in agricultural fields and in home-gardens. It belongs to the genus *Ocimum* which is comprised of up to 160 different species (Paton *et al.*, 1999). *O. basilicum* is characterized by a large diversity among its genotypes from various aroma types, leaf size and shape, leaf and stem color, inflorescence color and structure, grow habit and seeds morphology (Dudai *et al.*, 2018). These diverse genotypes have a wide range of uses and proposes, mainly as a culinary herb but also for their essential oil, as a fragrant for the cosmetics industry, as a source for health beneficial molecules such as antioxidants, as well as for ornamental proposes. It is widely used around the world in various cuisines from Italy to Thailand and from United States to Columbia. Mainly appreciated for its unique aroma, accumulated in specialized glandular trichomes, in every kitchen a different basil genotype, carrying a different aroma profile, is preferred. Sweet basil, the type of basil used in the famous Italian Pesto sauce, is the most widespread basil grown in the world. Basil breeding programs leg behind other crops, such as wheat, rice or tomato, due to lack of genomic data and unclear chromosome number and ploidy level.

The genome size, chromosomes number and ploidy levels of various basil genotypes were estimated using flow cytometry and chromosomes count methods. The results, especially regarding chromosomes number varied greatly between studies and between genotypes. For *O. basilicum* chromosomes number, 2n value were found to be 48, 52, 50, 52, 56, 72 or 74 (Pushpangadan and Sobti, 1982, Paton and Putievsky, 1996, Paton *et al.*, 1999, Mukherjee *et al.*, 2005). More recently, Carović-Stanko *et al.* (2010) found 2n = 48 for 20 *O. basilicum* genotypes and 2n = 72 for two *O. basilicum* var. *purpurascens* Benth. Also, for other species of the genus *Ocimum*, the chromosome number was found to vary greatly between and within species. For the holy basil, *O. tenuiflorum* L. (*O. sanctum*), 2n was found to be 16, 32, 36 and 72 (Paton and Putievsky, 1996, Mukherjee and Datta, 2005, Rastogi *et al.*, 2014). Several works estimated the size of the genome of *O. basilicum* based on DNA content measurements. With the rough assumption that 1 pg DNA = 0.978 Mbp (Dolezel *et al.*, 2003), the haplotypic (1C) genome size of 28 *O. basilicum* genotypes varied from 1.46 Gbp to 2.32 Gbp (Carović-Stanko *et al.*, 2010, Koroch *et al.*, 2010). For the cultivar ‘Perrie’, a the genome size was estimated to be 1.56 Gbp (Koroch *et al.*, 2010). The genome size of the holy basil was estimated to be 1.39 Gbp in one study (Koroch *et al.*, 2010) and 0.35 Gbp in another study (Carović-Stanko *et al.*, 2010). Recently, two works sequenced the genome of the holy basil and assembled 386 Mbp (Rastogi *et al.*, 2015) and 374 Mbp (Upadhyay *et al.*, 2015). The scaffolds N50 in these two studies were 303 kbp (Rastogi *et al.*, 2015) and 27 kbp (Upadhyay *et al.*, 2015) with total of 9,059 and 78,224 scaffolds, respectively.

While being cultivated worldwide, a reference genome of *O. basilicum* was not published so far. Actually, very limited next-generation sequencing (NGS) data of basil were published. Almost two decades ago, comprehensive EST libraries of several genotypes were generated and several works characterizing genes of aroma biosynthesis pathways were published (Gang *et al.*, 2001, Iijima *et al.*, 2004). More recently, a transcriptome of *O. basilicum*, together with a transcriptome of *O. tenuiflorum*, were generated using RNA-sequencing finding 130,043 and 69,117 transcripts, respectively (Rastogi *et al.*, 2014). Later, genotyping-by-sequencing approach (ddRAD-seq) was applied to an F_2_ mapping population derived from a cross between two *O. basilicum* genotypes to generate a linkage genetic map. The map span over 26 linkage groups with a total genetic length of 3031 cM (Pyne *et al.*, 2017). Yet, the lack of a reference genome for *O. basilicum* prevented an effective utilization of next-generation sequencing data. Here we report for the first time on the sequence of *O. basilicum* high-quality genome based on paired-end, mate-pair and Chromium^TM^ libraries. It was found to that the haplotype genome size of the cultivar ‘Perrie’ is 2.16 Gbp comprised of 12,212 scaffolds with N50 greater than 19Mbp with only 107 scaffolds comprising more than 90% of the total assembly. With an established reference genome, scientists will be able to better design NGS studies and make more knowledge-based decisions.

## Materials and methods

### Plant material

We used the cultivar ‘Perrie’ that is a sweet basil with inverted spoon-like shape leaves and typical aroma of “Genovese” basil (Figure 1).

**Figure 1.**
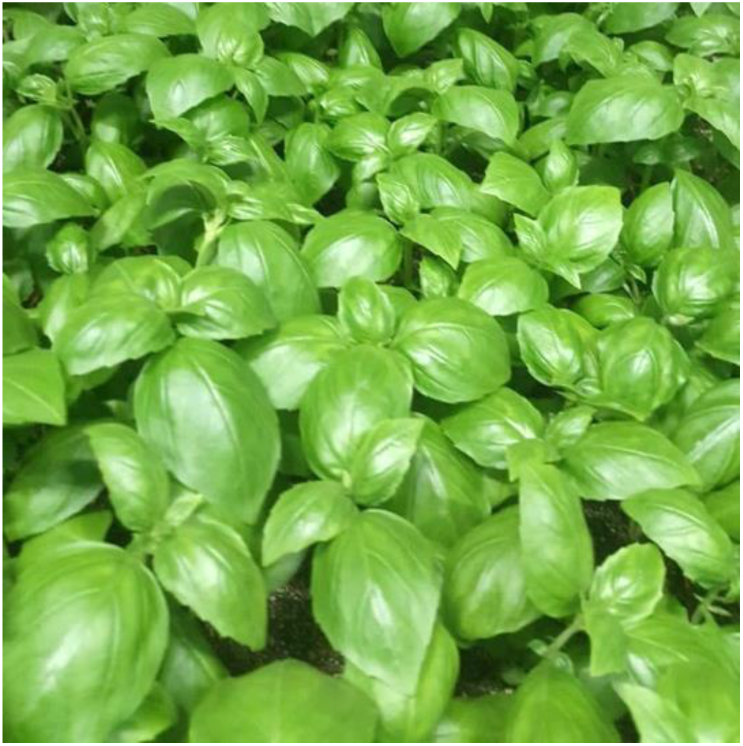
Basil of the cultivar ‘Perrie’

### Genomic DNA extraction

For high molecular weight DNA extraction, 150mg etiolated basil leaves were harvested, frozen in liquid nitrogen and ground with mortar and pestle. We followed Healey *et al.* (2014) with slight modifications. The fine powder was transferred to 15 ml tube and 1.5 ml of extraction buffer (100 mM Tris pH7.5, 1.5 M NaCl, 20 mM EDTA, 2% w/v CTAB, 2% w/v PVP-40, 0.3% v/v β-mercaptoethanol) was added and the samples were incubated at 60°C for 1 h with occasional shaking. Samples were centrifuged for 5 min at 5000 *g* in 4°C, and the supernatant was transferred to a new tube. One volume of isoamyl alcohol: chloroform (1:24 v/v) was added and samples were gently mix for 5 minutes. Then, samples were centrifuged for 10 min at 5000 *g* in 4°C and the supernatant was transferred to a new tube. RNAse was added to final concentration of 20 µg / ml and samples were incubated for 15 min at 37°C with occasional shaking. Then, 1 volume of isoamyl alcohol: chloroform (1:24 v/v) was added and samples were gently mixed for 5 min followed by 10 min centrifugation at 5000 *g* in 4°C. The supernatant was transferred to new tube and half volume of 5 M NaCl was added and samples were gently mixed. Then, 3 volumes of ice-cold 95% ethanol were added and samples were mixed for 2 min following incubation at −20°C for 1 h. Samples were centrifuged for 10 min at 5000 *g* in 4°C and the supernatant was discarded and samples were washed with 3 ml of ice-cold 70% ethanol. Finally, samples were centrifuged for 10 min at 5000 *g* in 4°C, the ethanol was discarded and tubed were left open to evaporate ethanol residues. Then, the DNA was resuspended in 75 µl of 37°C TE buffer. DNA quality and quantity were evaluated with nanodrop spectrophotometer and Qubit fluorometer and by running overnight on a 0.5% agarose gel.

### Library construction and DNA sequencing

Five size-selected genomic DNA libraries ranging from 470bp to 10Kb were constructed for each material. Two shotgun Paired-End (PE) libraries were made using DNA template fragments size selected of ~470bp with no PCR amplification (PCR-free). This fragment size was designed to produce a sequencing overlap of the fragments to be sequenced on the Hiseq2500 v2 Rapid run mode as 2×265bp, thus creating an opportunity to produce ‘stitched’ reads of approximately 265bp to 520bp in length. Two genomic libraries of 700bp DNA fragment sizes were prepared using the TruSeq DNA Sample Preparation Kit version 2 with no PCR amplification (PCR-free) according to the manufacturer’s protocol (Illumina, San Diego, CA). Three separate Mate-Pair (MP) libraries were constructed with 2-5Kb, 5-7Kb and 8-10Kb jumps using the Illumina Nextera MP Sample Preparation Kit (Illumina, San Diego, CA). The 700bp shotgun libraries and all 3 MP libraries were sequenced on Illumina NovaSeq6000 S4 flowcell as 2×150bp reads. In addition, High molecular weight DNA was prepared and the quality of the DNA samples was verified by pulsed-field gel electrophoresis. The DNA fragment longer than 50Kb was subjected to construct one Gemcode library using the Chromium instrument (10X Genomics, Pleasanton, CA). This library was sequenced on NovaSeq6000 S4 flowcell as 2×150bp reads. Sequencing strategy is detailed in Table 1.0. Paired-end, MP and Chromium libraries construction and sequencing were conducted at Roy J. Carver Biotechnology Center, University of Illinois at Urbana-Champaign.

**Table 1.** Contigs statistics summary

|  |  |
| --- | --- |
| Total contigs | 128,921 |
| Assembly size | 2,105,853,635 |
| N50 | 45,710 |
| N50 #sequences* | 12,028 |
| N90 | 8,583 |
| N90 #sequences** | 53,359 |
| MAX | 585,067 |
| *-number of contigs comprising 50% of the total length |  |
| **-number of contigs comprising 90% of the total length |  |

### Genome Assembly

Genome assembly was conducted using DeNovoMAGIC™ software platform (NRGene Ltd., Ness Ziona, Israel). This is a DeBruijn-graph-based assembler, designed to efficiently extract the underlying information in the raw reads to solve the complexity of the DeBruijn graph due to genome polyploidy, heterozygosity and repetitiveness. This task is accomplished using accurate-reads-based traveling in the graph that iteratively connected consecutive phased contigs over local repeats to generate long phased scaffolds (Lu *et al.*, 2015, Hirsch *et al.*, 2016, Avni *et al.*, 2017, Luo *et al.*, 2017, Zhao *et al.*, 2017). Assembly results are summarized in table 2.0. In brief, the algorithm is composed of the following steps:

**Table 2.** Scaffolds statistics summary

|  |  |
| --- | --- |
| Total scaffolds | 12,212 |
| Assembly size | 2,133,958,912 |
| Gaps size | 27,347,277 |
| Gaps % | 1.28 |
| N50 | 19,298,043 |
| N50 #sequences* | 33 |
| N90 | 5,853,927 |
| N90 #sequences** | 107 |
| MAX | 71,596,212 |
| *-number of scaffolds comprising 50% of the total length |  |
| **-number of scaffolds comprising 90% of the total length |  |

1. Reads pre-processing. PCR duplicates, Illumina adaptor AGATCGGAAGAGC and Nextera linkers (for MP libraries) were removed. The PE 450bp 2×265bp libraries overlapping reads were merged with minimal required overlap of 10bp to create the stitched reads.
2. Error correction following pre-processing, merged PE reads were scanned to detect and filter reads with putative sequencing error (contain a sub-sequence that does not reappear several times in other reads).
3. Contigs assembly. The first step of the assembly consists of building a DeBruijn graph (kmer=127 bp) of contigs from the all PE and MP reads. Next, PE reads were used to find reliable paths in the graph between contigs for repeat resolving and contigs extension.
4. Scaffolds assembly. Later, contigs were linked into scaffolds with PE and MP information, estimating gaps between the contigs according to the distance of PE and MP links.
5. Fill Gaps. A final fill gap step used PE and MP links and DeBruijn graph information to detect a unique path connecting the gap edges.
6. Scaffolds elongation and refinement. The 10X barcoded reads were mapped to the assembled scaffolds and clusters of reads with the same barcode mapped to adjacent contigs in the scaffolds were identified to be part of a single long molecule. Next, each scaffold was scanned with a 20kb length window to ensure that the number of distinct clusters that cover the entire window (indicating a support for this 20kb connection by several long molecules) was statistically significant with respect to the number of clusters that span the left and the right edge of the window. In case where a potential scaffold assembly error was detected the scaffold was broken at the two edges of the suspicious 20kb window. Finally, the barcodes that were mapped to the scaffold edges were compared (first and last 20kb sequences) to generate a scaffolds graph with a link connecting two scaffolds with more than two common barcodes. Linear scaffolds paths in the scaffolds graph were composed into the final scaffolds output of the assembly.

### Essential oil extraction and analysis

Fresh leaves and stems (300g) were distilled according to Putievsky *et al.* (1999) by a modified Clevenger type apparatus, and analyzed using a gas-chromatograph mass-detector (Agilent, www.agilent.com/) according to Dudai *et al.* (2003).

## Results and discussion

### Genome sequencing and assembly

To sequence and assemble the genome of *O. basilicum*, we have extracted high-molecular weight DNA form etiolated cotyledons of the cultivar ‘Perrie’ which is a very stable FOB resistant line (Figure 2). DNA libraries were prepared and sequenced with Illumina instruments as described in material and methods. We generated a couple of PCR-free paired-end libraries (insert sizes of ~ 450 bp and of ~ 750 bp) which produced 136 Gbp data. In addition, we have generated three mate-pair libraries with different insert sizes (2-4 Kbp, 5-7 Kbp and 8-10Kbp) which produced 98 Gbp data. Finally, we use 10Xgenomics^TM^ technology to add 23 Gbp of third generation sequencing data. Altogether, we generated ~ 257 Gbp of short reads data. Based on the previous estimation of the genome size of ‘Perrie’ cultivar as 1.56 Gbp long (Koroch *et al.*, 2010), the data generated represent 165X genome coverage.

**Figure 2.**
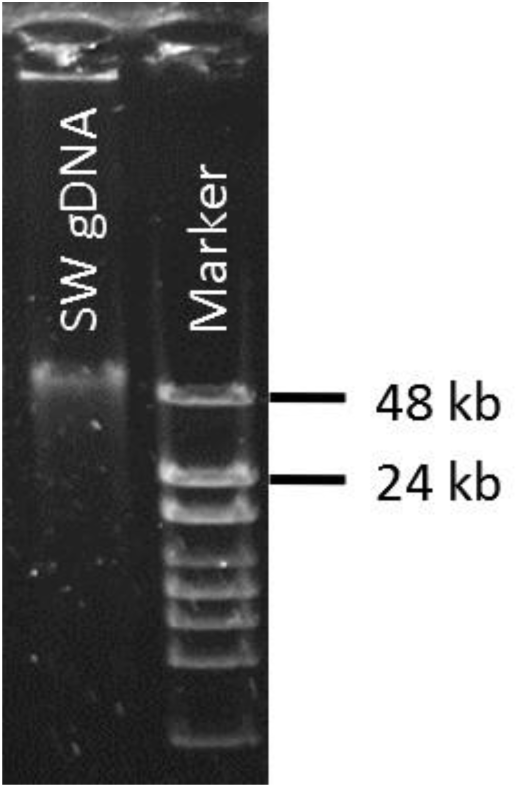
Gel capture of the genomic DNA extracted for the sequencing of the sweet basil genome. SW, sweet basil; gDNA, genomic DNA

To assemble the short reads information into contigs and scaffolds we used DeNovoMAGIC^TM^ assembler v. 3. We generated 128,921 contigs to a total size of 2.105 Gbp and N50 of 45,710 bp (Table 1). We then assembled the contigs into scaffolds resulted in a total genome size was ~2.13 Gbp comprised of 12,212 scaffolds (Table 2). The gaps size was ~ 27.3 Mbp, representing less than 1.3% of the total assembly size. The N50 value was approximately 19 Mbp and N90 value was approximately 5.8 Mbp. The largest scaffold was 71.6 Mbp and only 107 scaffolds comprising more than 90% of the assembled genome. Considering haplotypic chromosome number of n = 24, these last two parameters suggest some of the scaffolds are almost in a size of a complete chromosome indicating the high-quality of the assembly. The contigs number was 128,921 with N50 value of 47,710bp. Additionally, ~1.1 million sequences which couldn’t be placed within the assembly generated a total size of ~0.34 Gbp and N50 of 384 bp. The assembly suggested a very high level of homozygosity of the studied cultivar, ‘Perrie’. A size of 2.13 Gbp for the haplotype genome of ‘Perrie’ cultivar is quite larger from the 1.56 Gbp estimated by cytological measurements of DNA content found by (Koroch *et al.*, 2010). Yet, it is similar to the values of other 20 cultivars of *O. basilicum* found by (Carović-Stanko *et al.*, 2010) that ranged from 2.04 to 2.32 Gbp. The assembly size of 2.13 Gbp still embodies a very high genome coverage of 121X.

**Table 3.** BUSCO statistics summary

|  |  |
| --- | --- |
| Complete BUSCO genes | 1339 (93.0%) |
| Complete BUSCO genes - Single-Copy | 267 (18.5%) |
| Complete BUSCO genes - Duplicated | 1072 (74.4%) |
| Fragmented BUSCO genes | 21 (1.5%) |
| Missing BUSCO genes | 80 (5.6%) |
| Total BUSCO genes searched | 1440 |

**Table 4.**
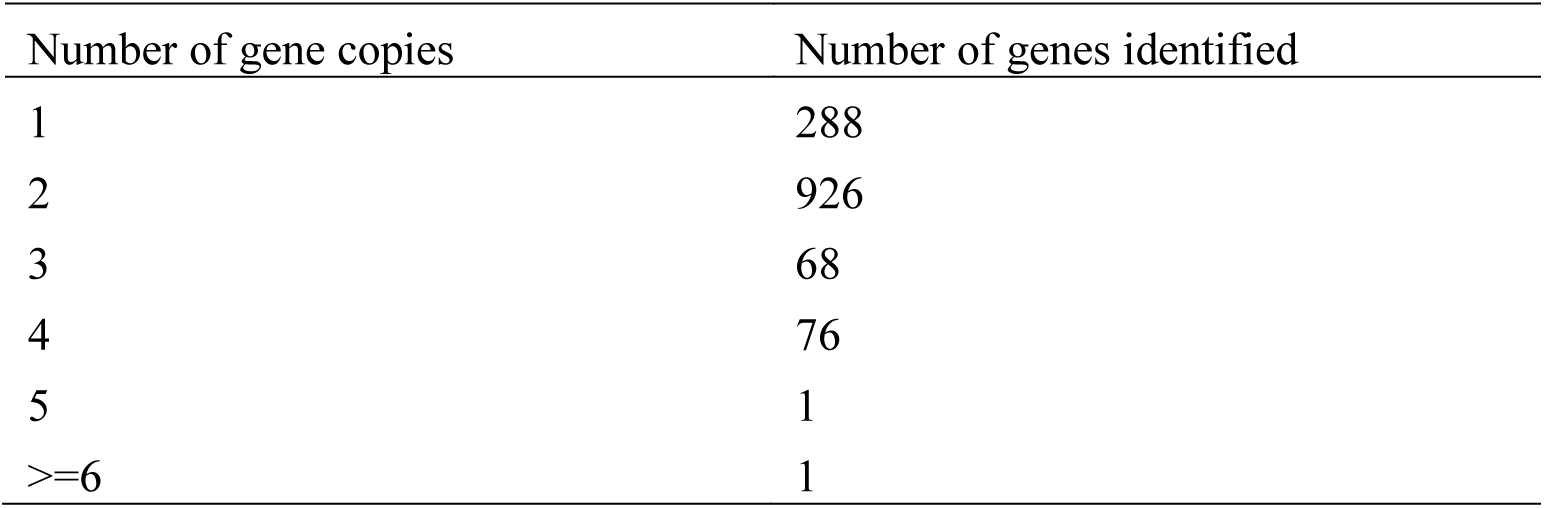
BUSCO copy number analysis summary

To estimate the quality of our assembly we performed an independent analysis with the BUSCO pipeline which is comprised of a set of 1440 single-copy ortholog genes (Simao *et al.*, 2015). Of this set, 93% of the genes were found to be complete, 1.5% were found to be fragmented and 5.6% were missing. An interesting finding was that 74.4% of the BUSCO gene were found in multicopy state with most of them (86%) being duplicated. This indicates that *O. basilicum* is a tetraploid species as was suggested before (Pyne *et al.*, 2017).

### Phenotypic characteristics of ‘Perrie’ cultivar

The cultivar that was used for the sequencing of the sweet basil reference genome, ‘Perrie’, is a well-established Fusarium wilt resistant cultivar (Dudai *et al.*, 2002). It is the chief cultivar grown in Israel for year round production of fresh-cut herb. We present here a full description of the essential oil composition of ‘Perrie’ cultivar grown in greenhouses as gold-standard for high-quality aroma of fresh sweet basil (Table 5). The essential oil was content was 0.13% of the fresh weight and composed of monoterpenes (45.4%), sesquiterpenes (26.2%), phenylpropanoids (27.6%) and traces of fatty acid-derived straight-chain volatiles (0.1%). The main compounds were the monoterpenes linalool (30.5%) and 1,8-cineole (9.5%), and the phenylpropanoid eugenol (27.4%). These were followed by the sesquiterpenes τ-cadinol, germacrene D, γ-cadinene and bicyclogermacrene (total of 14.3%). In total, 62 compounds were detected including 33 sesquiterpenes, 21 monoterpenes, two phenylpropanoids and six fatty acid-derived compounds. The aroma profile of ‘Perrie’ is clean of the anise-like aroma compound methyl chavicol which is the main compound in Thai basil (Dudai and Belanger, 2016). Many breeding efforts for various traits such as disease resistances and yield fail to eliminate the presence of methyl chavicol preventing the marketing of the product as ‘Genovese’ basil due to variable levels of anise-like aroma. Now, with genome sequence in hand, application of molecular breeding tools will make such efforts more precise, effective and in a timely manner. It will facilitate QTL studies to better understand underlying mechanisms controlling tolerance to multiple environments and the biochemistry and chemistry of secondary metabolite production. Ultimately, it will bring about employment of genomic selection in basil breeding.

**Table 5.**
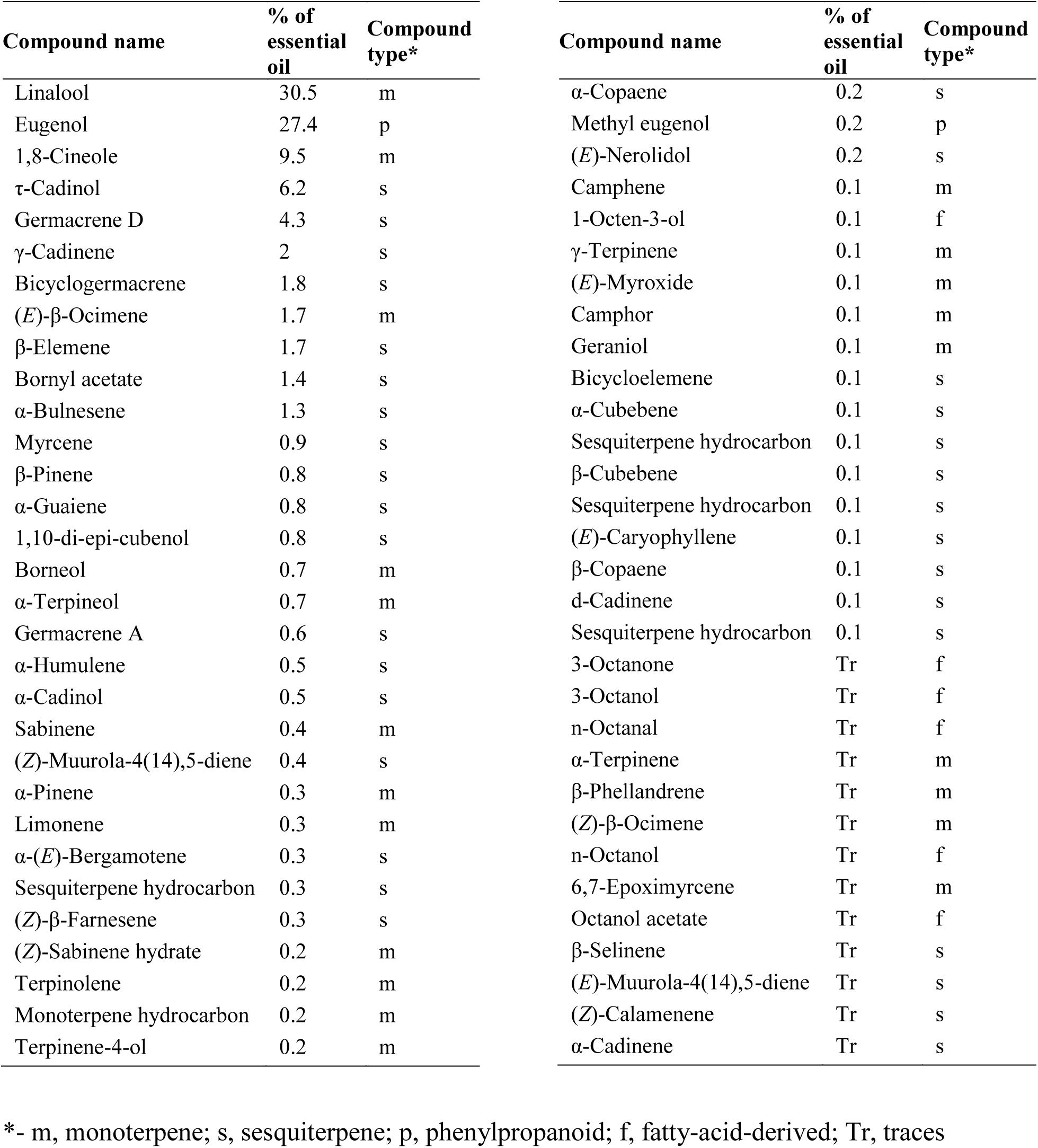
Essential oil composition of sweet basil from ‘Perrie” cultivar

| Compound name | % of essential oil | Compound type* | Compound name | % of essential oil | Compound type* |
| --- | --- | --- | --- | --- | --- |
| Linalool | 30.5 | m | α-Copaene | 0.2 | s |
| Eugenol | 27.4 | p | Methyl eugenol | 0.2 | p |
| 1,8-Cineole | 9.5 | m | (*E*)-Nerolidol | 0.2 | s |
| τ-Cadinol | 6.2 | s | Camphene | 0.1 | m |
| Germacrene D | 4.3 | s | 1-Octen-3-ol | 0.1 | f |
| γ-Cadinene | 2 | s | γ-Terpinene | 0.1 | m |
| Bicyclogermacrene | 1.8 | s | (*E*)-Myroxide | 0.1 | m |
| (*E*)-β-Ocimene | 1.7 | m | Camphor | 0.1 | m |
| β-Elemene | 1.7 | s | Geraniol | 0.1 | m |
| Bornyl acetate | 1.4 | s | Bicycloelemene | 0.1 | s |
| α-Bulnesene | 1.3 | s | α-Cubebene | 0.1 | s |
| Myrcene | 0.9 | s | Sesquiterpene hydrocarbon | 0.1 | s |
| β-Pinene | 0.8 | s | β-Cubebene | 0.1 | s |
| α-Guaiene | 0.8 | s | Sesquiterpene hydrocarbon | 0.1 | s |
| 1,10-di-epi-cubenol | 0.8 | s | (*E*)-Caryophyllene | 0.1 | s |
| Borneol | 0.7 | m | β-Copaene | 0.1 | s |
| α-Terpineol | 0.7 | m | d-Cadinene | 0.1 | s |
| Germacrene A | 0.6 | s | Sesquiterpene hydrocarbon | 0.1 | s |
| α-Humulene | 0.5 | s | 3-Octanone | Tr | f |
| α-Cadinol | 0.5 | s | 3-Octanol | Tr | f |
| Sabinene | 0.4 | m | n-Octanal | Tr | f |
| (*Z*)-Muurolo-4(14),5-diene | 0.4 | s | α-Terpinene | Tr | m |
| α-Pinene | 0.3 | m | β-Phellandrene | Tr | m |
| Limonene | 0.3 | m | (*Z*)-β-Ocimene | Tr | m |
| α-(*E*)-Bergamotene | 0.3 | s | n-Octanol | Tr | f |
| Sesquiterpene hydrocarbon | 0.3 | s | 6,7-Epoximycene | Tr | m |
| (*Z*)-β-Farnesene | 0.3 | s | Octanol acetate | Tr | f |
| (*Z*)-Sabinene hydrate | 0.2 | m | β-Selinene | Tr | s |
| Terpinolene | 0.2 | m | (*E*)-Muurolo-4(14),5-diene | Tr | s |
| Monoterpene hydrocarbon | 0.2 | m | (*Z*)-Calamenene | Tr | s |
| Terpinene-4-ol | 0.2 | m | α-Cadinene | Tr | s |

\*- m, monoterpene; s, sesquiterpene; p, phenylpropanoid; f, fatty-acid-derived; Tr, traces

